# Cross-Species Experiments Reveal Widespread Cochlear Neural Damage in Normal Hearing

**DOI:** 10.1101/2021.03.17.435900

**Authors:** Hari M. Bharadwaj, Alexandra R. Hustedt-Mai, Hannah M. Ginsberg, Kelsey M. Dougherty, Vijaya Prakash Krishnan Muthaiah, Anna Hagedorn, Jennifer M. Simpson, Michael G. Heinz

## Abstract

Animal models suggest that cochlear afferent nerve endings may be more vulnerable than sensory hair cells to damage from acoustic overexposure and aging, but that such damage cannot be detected in standard clinical audiometry. Co-ordinated experiments in at-risk humans and a chinchilla model using two distinct physiological assays suggest that cochlear neural damage exists even in populations without clinically recognized hearing loss.

## Introduction

Acoustic overexposure and aging are conventionally thought to cause hearing damage by damaging sensory cells (hair cells) in the cochlea leading to a decrease in hearing sensitivity (Rabinowitz et al., 2006; Cruickshanks et al., 2010). Such loss of sensitivity is quantified using threshold audiometry, which is the foundation of current clinical diagnostics, audiological counseling, and patient management. Contrary to this view, recent animal data show substantial permanent damage to synapses and auditory afferents innervating the cochlea from noise exposure (NE; Kujawa and Liberman, 2015), even in the absence of hair-cell damage. In normal aging, such primary cochlear neural degeneration is evident not only in animal models (Sergeyenko et al., 2013), but also in analyses of post-mortem human temporal bones (Wu et al., 2019). Insidiously, even an extreme degree of such cochlear synaptopathy (CS) is unlikely to affect thresholds on clinical audiograms (Lobarinas et al., 2013). Thus, the extent to which such “hidden” damage occurs in behaving humans and contributes to suprathreshold perceptual deficits (e.g., listening in restaurants) is unknown and hotly debated.

Given that most patients seeking audiological help struggle in listening environments with background noise, considerable effort has been directed to examine associations between risk factors for CS (i.e., NE history, or age), and suprathreshold hearing in noise. Such investigations have yielded mixed results (Bharadwaj et al., 2015; Prendergast et al., 2017; Bramhall et al., 2017; Yeend et al., 2017; Grant et al., 2020), perhaps because they have been hampered by multiple sources of variability (Bharadwaj et al., 2019). First, individuals with a history of greater NE and more advanced age tend to also have greater audiometric threshold elevations making it difficult to accurately estimate the effects attributable to CS. Second, NE history and the history of hearing protection use are difficult to assess retroactively; moreover, cumulative NE levels tend to correlate with age rendering it difficult to disentangle their respective effects. Finally, while non-invasive assays correlated with CS have been successful in certain mouse strains (Furman et al., 2013; Valero et al., 2018), whether such assays are sensitive when there are genetic and experiential factors introducing considerable extraneous variance is yet to be established. In the present study, many of these impediments were mitigated by (1) employing two distinct non-invasive assays that were specifically designed to reduce the variability from commonly occurring extraneous factors (Bharadwaj et al., 2019), and (2) using *parallel measurements in two species*, a “wild-type” chinchilla model of cochlear synaptopathy with minimal hair-cell loss (Hickox et al., 2017), and human groups with *substantially different risk for synaptopathy but tightly matched clinical audiograms*.

## Results and Discussion

To examine the integrity of the afferent cochlear nerve, two primary pathways driven by the same auditory afferents were evaluated. The first assay measured the auditory brainstem response (ABR) wave-I amplitude in response to high-pass (HP) clicks (3–8 kHz, for humans) or high-frequency tone-bursts (4 and 8 kHz, for chinchillas), normalized by the wave-V amplitude. The human ABR was acquired using ear-canal “tiptrodes” and 32-channel scalp measurements. The second assay was the wideband middle-ear muscle reflex (WB-MEMR) in response to broadband (BB, 0.5–8 kHz, both species) and HP (3–8 kHz, humans only) noise elicitors. These specific choices were guided by a series of prior experiments, and help reduce extraneous variance from cochlear dispersion (including sex effects), head/brain anatomy, ear-canal filtering effects, and spectral-profile variations in middle-ear reflex properties (Bharadwaj et al., 2019). In addition to these putative CS assays, distortion-product otoacoustic emissions (DPOAEs) were obtained with f2-primaries spanning 2–16 kHz to control for outer-hair-cell (OHC) damage. Behavioral audiometry from 250 Hz–16 kHz was also done in humans.

To establish the sensitivity of our assays in a genetically heterogeneous cohort, but with known lab-induced pathophysiology, we studied chinchillas exposed to moderate-level noise in a pre-post design (Fig. 1A). Awake chinchillas were exposed to 100-dB-SPL octave-band noise centered at 1 kHz for two hours, which produces histologically confirmed CS with no significant permanent threshold shift (Hickox et al., 2017, Fig. 1B). The CS profile spans 1–10 kHz cochlear places, similar to findings in post-mortem temporal bones of middle-aged humans (Wu et al., 2019). Accordingly, DPOAEs are reduced in magnitude one-day post-NE but recover to pre-exposure levels two-weeks post-NE consistent with temporary threshold shifts (TTS) but no permanent damage to OHCs (Fig. 1C). ABR threshold shifts at two-weeks post-NE are < 5 dB (Fig. 1D). In contrast, the WB-MEMR (Fig. 1E) showed a large (> 50%) reduction in amplitudes and elevation in thresholds that did not recover even two-weeks post-NE (F(11, 120) = 6.74, P = 1e-8, N=7). This suggests that the WB-MEMR is highly sensitive to CS even in a genetically heterogeneous cohort of animals despite likely including many extraneous sources of variance. As a second assay, the ratio of the ABR wave-I-to-wave-V ratio was computed for high-amplitude 4-kHz and 8-kHz tone bursts to mirror our human HP click protocol (Bharadwaj et al., 2019). This assay also showed a reduction post-NE that did not recover at two weeks, but not to a statistically significant degree by conventional criteria (Fig. 1F). Taken together, these results show that while both assays may be sensitive to CS, the WB-MEMR shows greater promise in the presence of genetic heterogeneity.

**Figure 1.**
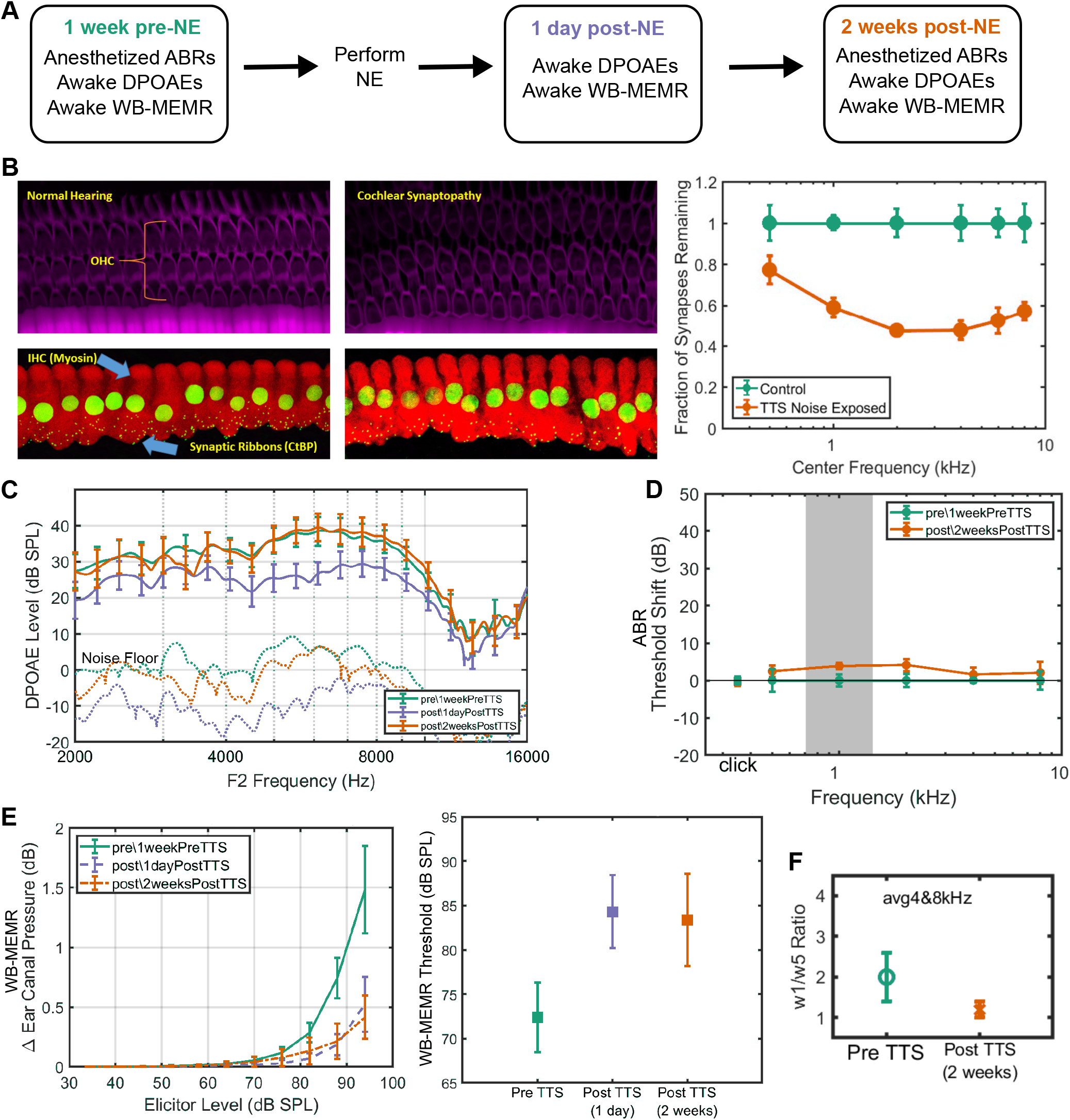
Exposing chinchillas to octave-band noise causes TTS and transient DPOAE reductions but sustained CS, reduced WB-MEMR and suprathreshold ABR. (A) Exposure and measurement timeline. (B) Confocal imaging reveals broad CS following TTS exposure. (C) Reduced DPOAE amplitudes 1-day post-NE, but full recovery by 2 weeks. (D) ABR thresholds at 2-weeks post-NE are within 5 dB of pre-NE levels. (E) Large (>50%) sustained reduction in WB-MEMR amplitudes and increased thresholds. (F) Reduced ABR wave-I/V ratio at 2-weeks post. All datapoints are estimated mean ± STE bars.

While it is well established that multiple rodent models are susceptible to CS, whether humans show the same vulnerabilities is still contested (Dobie and Humes, 2017; Bramhall et al., 2019). To investigate whether behaving humans show evidence of CS, we studied three groups with varying risk in a cross-sectional design (N=166 individuals total). The control group (“YCtrl”, N=55) comprised of young (18—35 years of age) individuals. A middle-aged cohort (“MA”, 36–60 years of age, N=58) formed our first high-risk group, by virtue of their age. A second high-risk cohort (N=53) of young individuals (aged 18–35 years) with regular and substantial acoustic exposures (“YExp”) was recruited from the Purdue University marching band and shooting clubs. All groups had clinically normal audiograms (thresholds better than 25 dB HL up to 8 kHz), and crucially, were matched (Fig. S-1A) in both mean and median threshold within <5 dB at every audiometric frequency in the standard clinical range (0.25–8 kHz). Furthermore, the YCtrl and the YExp groups were matched in mean and median thresholds (<5 dB) even in the extended-high-frequency range (8–16 kHz). This design helps dissociate effects of audiometric loss and the effects of cochlear synaptopathy. In a conservative addition to this design, the effects of residual individual variation in audiometric thresholds were explicitly accounted for during analysis through a linear mixed-effects modeling approach.

Consistent with the audiometric data, DPOAE amplitudes were similar across the three groups in the clinical frequency range with the MA group showing some reductions near and beyond 8 kHz (Fig. 2A). Despite groups being tightly matched in clinical hearing status, the WB-MEMR data revealed significant group differences. With the BB noise elicitor (0.5–8 kHz), the MA group showed significantly reduced MEMR growth functions compared to the YCtrl group when controlling either for the audiometric variations (F(10, 939) = 2.7, P = 0.0028) or for DPOAE amplitudes (F(10, 941) = 2.8, P = 0.0020), whereas the YExp group did not (Fig. 2B). With the HP noise elicitor (3–8 kHz), both high-risk groups showed significantly attenuated MEMR growth function (Fig. 2D), when controlling for either audiometric variations (YCtrl vs. YExp: F(10, 896) = 3.71, P = 7e-5; YCtrl vs. MA: F(10, 939) = 8.57, P = 1.7e-13), or for DPOAE amplitudes (YCtrl vs. YExp: F(10, 898) = 3.84, P = 4.2e-5; YCtrl vs. MA: F(10, 941) = 8.91, P = 4.2e-14). These results suggest substantial CS in the MA group spanning a broad frequency range, similar to our chinchilla model and to previous indications from human post-mortem data (Wu et al., 2019). On the other hand, the results in the YExp group suggest a lesser degree of CS that is localized to the basal regions of the cochlea. These results are corroborated by the findings from our second assay where the ABR wave-I/V ratio for HP (3–8 kHz) clicks is attenuated in both high-risk groups, but more so in the MA group (Fig. 2C) after adjusting for audiometric variations (YCtrl vs. YExp: T(1, 155) = 1.89, P = 0.03; YCtrl vs. MA: T(1,155) = 3.794, P = 0.0001), or for DPOAE amplitudes (YCtrl vs. YExp: T(1, 156) = 1.731, P = 0.047; YCtrl vs. MA: T(1, 156) = 6.024, P = 5e-9). Consistent with the chinchilla findings, a comparison of the test statistics and P-values between WB-MEMR assays and ABR wave-I/V ratio suggests that the HP-noise-elicited WB-MEMR growth function is the most robust measure separating the control and high-risk human groups.

**Figure 2.**
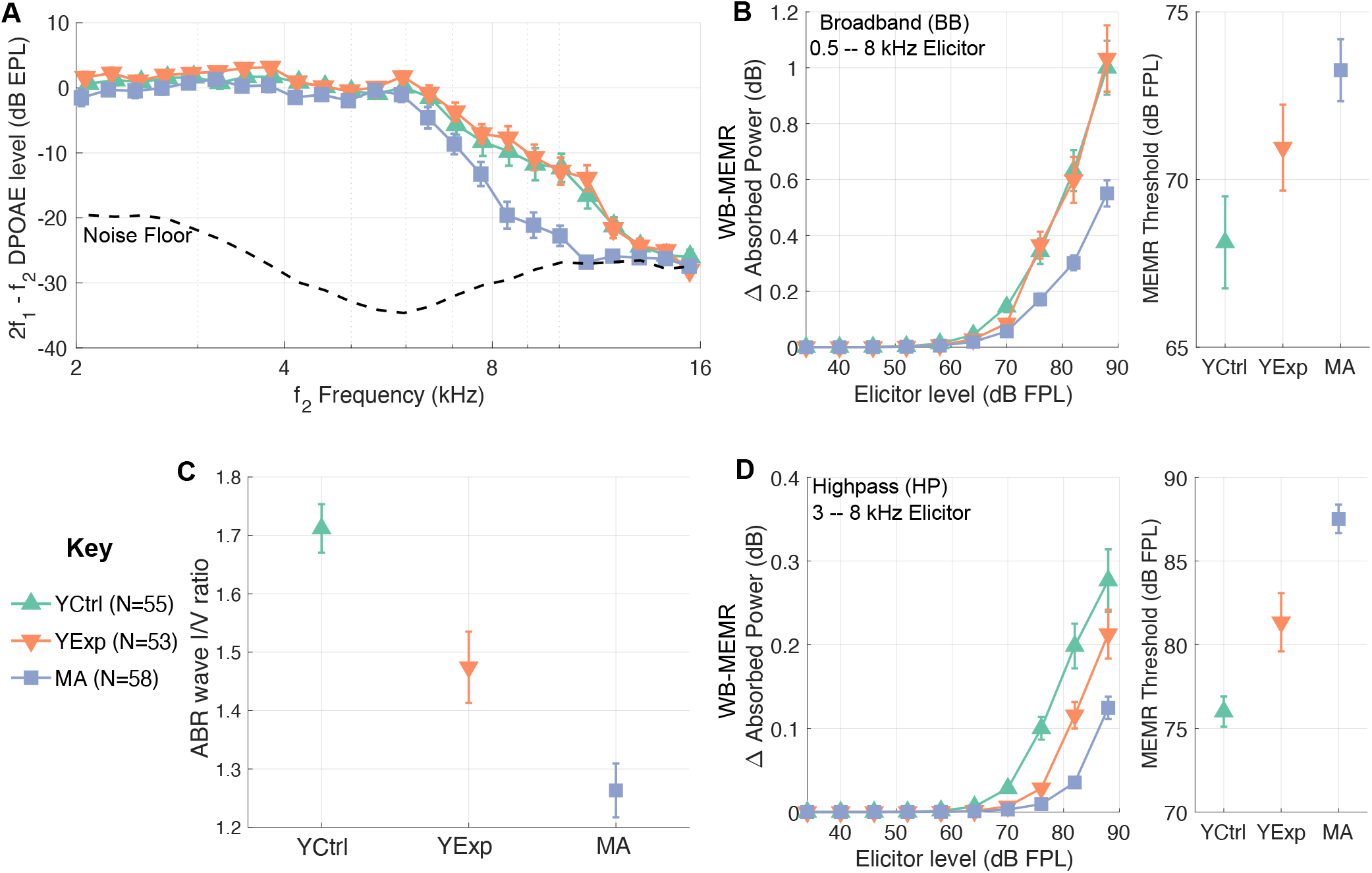
YExp and MA human groups show reduced WB-MEMR and ABR compared to YCtrl despite tightly matched audiograms. (A) DPOAE amplitudes are closely matched up to 8 kHz. The MA group shows reduced DPOAEs between 8 to 16 kHz, consistent with audiograms. WB-MEMR to BB elicitors is significantly reduced in the MA group (B), whereas both high-risk groups show reductions with HP elicitors (D). (C) ABR wave-I/V ratio is reduced in both high-risk groups. All datapoints are estimated mean ± STE bars.

To test the sensitivity of clinically available versions of the MEMR and ABR, the same human groups were also studied using standard clinical equipment and protocols. A comparison of raw effect sizes for various lab and clinic-style measures showed that the clinical measures, although less sensitive, yielded results consistent with our more targeted laboratory assays (Fig. 3A), corroborating the notion that the high-risk groups exhibit CS. Finally, to obtain further insight into the sensitivity of the clinical MEMR, we also analyzed data from a large publicly available bank of audiological measurements from the NHANES 2012 repository, which revealed a steady decline of the MEMR amplitude with age despite normal hearing sensitivity (Flamme et al., 2017, Fig. 3B).

**Figure 3.**
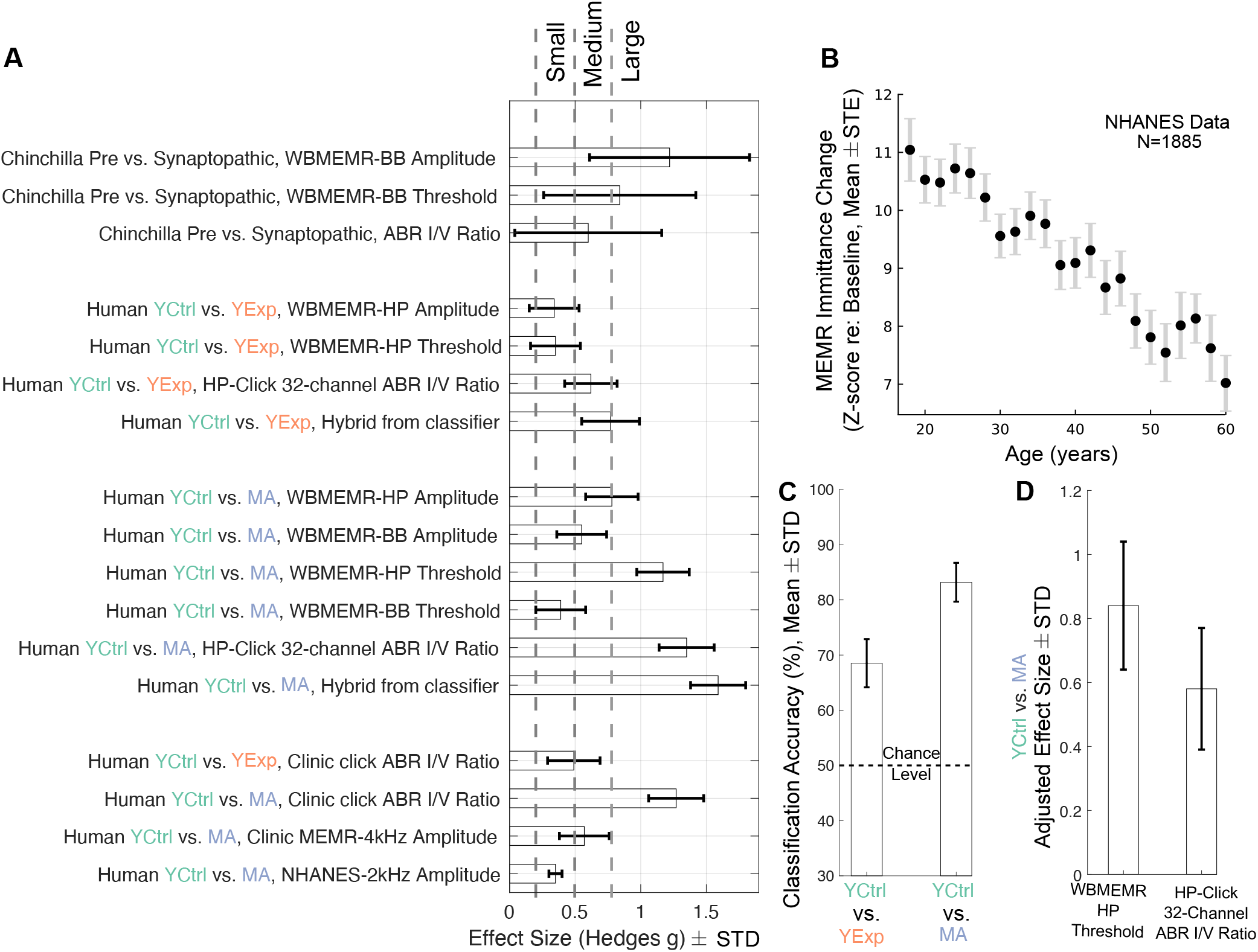
Clinic-style MEMR and ABR measures are also consistent with CS in high-risk human groups, but less sensitive than lab-style measures. (A) Effect-size comparisons for individual assays. (B) NHANES repository data show steady decline in MEMR amplitude with age despite normal audiograms. (C) An SVM classifier performs substantially above chance in blindly classifying individuals into their groups. (D) Effect size adjusted for audiometric thresholds *>* 8 kHz is larger for the “best” WB-MEMR than the “best” ABR metric.

A support-vector machine classifier was trained to use both WB-MEMR and ABR wave-I/V ratio metrics to blindly classify whether an individual belongs to the YCtrl group or one of the high-risk groups. Leave-one-out cross validation demonstrated substantially above-chance classification (Fig. 3C). Using conventional effect-size interpretations (Cohen, 1992), raw effect sizes in the small-medium range were obtained for individual physiological metrics for the YCtrl vs. YExp comparison, and medium-large for the combined hybrid metric extracted by the classifier. For YCtrl vs. MA, medium-large effect sizes were obtained for individual metrics and large effect size for the combined hybrid metric extracted by the classifier (Fig. 3A). Note that although raw effect sizes are larger for ABR than for MEMR metrics, the ABR wave-I amplitude is known to also be strongly correlated with audiometric thresholds and DPOAE amplitudes in the extended-high-frequency range, which complicates the interpretation of ABR-based metrics (Bharadwaj et al., 2019). Because individuals in the MA group showed elevated audiometric thresholds in the extended-high-frequency range compared to those in the YCtrl group, we used a simple linear regression to partially adjust for this effect from the “best” WB-MEMR and the “best” ABR metrics. When effect sizes were re-estimated from the adjusted metrics, WB-MEMR thresholds (with HP elicitors) still showed a large effect size whereas the ABR wave-I/V effect size dropped to the medium range (Fig. 3D). Thus, when considering the chinchilla and human data together, WB-MEMR measures seem to be the most sensitive to subclinical cochlear-nerve damage, and also simpler to interpret in the face of audiometric loss beyond 8 kHz.

Taken together, our results suggest that humans are also susceptible to CS from NE and aging, and that such damage may be widespread even among individuals with good hearing status per current clinical criteria. Furthermore, the large (> 50%) reduction in the WB-MEMR amplitudes in our chinchillas and the parallel effects observed in at-risk human groups suggest that the WB-MEMR, particularly with a HP elicitor, is highly sensitive to CS. Our results support the interpretation that CS mediates the observed associations between WB-MEMR measures and speech-in-noise scores (Shehorn et al., 2020; Mepani et al., 2020). Lower sensitivity could also account for mixed results in similar comparisons using clinic-style MEMR measures (Mepani et al., 2020; Guest et al., 2019). The WB-MEMR assay is likely to be available for clinical use widely in the near future, and may also be appropriate as an outcome measure in clinical trials of therapeutic drugs intended to re-establish afferent cochlear synapses (Suzuki et al., 2016).

## Acknowledgments

Funding was provided by NIH R01DC015989 (HMB) and R01DC009838 (MGH), as well as the Purdue Summer Undergraduate Research Fellowship (SURF) Program (HMG). We would also like to thank Jay Gephart (Director, Purdue Bands) for assistance with subject recruitment, Brooke Zirlow and Satyabrata Parida for assistance with data collection, and Karolina Charaziak for input and sharing seed software code for FPL and EPL calibrations.

## Author Contributions

HMB, MGH, and JMS designed the study. ARH, HMG, KMD, and AH collected the data. VPKM performed histology. ARH and HMB analyzed the human data. MGH, HMG and HMB analyzed the chinchilla data. ARH, MGH, and HMB wrote the manuscript. All authors provided comments on the manuscript.

## Data Availability Statement

Individual human subject WB-MEMR responses for two different elicitors as a function of elicitor level, ABR wave I and V amplitudes, audiometric thresholds binned in three different frequency ranges, DPOAE amplitudes in the 3-8 kHz range and the 8-16 kHz range, age and gender can be obtained from https://figshare.com/^1^. Similarly, WB-MEMR, ABR, and OAE data for individual chinchillas at the pre-and 2-week-post-NE time points can be obtained from the same repository. The NHANES 2011-2012 audiological data is publicly available at https://www.cdc.gov/nchs/nhanes/

## Code Availability Statement

Our custom software for acquisition and analysis of electrophysiological responses, and for acquisition and analysis of acoustic responses (e.g., WB-MEMR, DPOAE) are publicly available at https://github.com/SNAPsoftware/ANLffr and https://github.com/SNAPsoftware/SNAPacoustics, respectively. The license is highly permissive; investigators interested in replicating the measures used in this study can adapt the code to match the specifics of their hardware. The same software was adapted for use with the chinchilla setup.

## Materials and Methods

### Human Groups

All human subject measures were conducted in accordance with protocols approved by the Purdue University Internal Review Board and the Human Research Protection Program. Participants were recruited via posted flyers and bulletin-board advertisements and provided informed consent. All participants had thresholds of 25 dBHL or better at audiometric frequencies in the 0.25–8 kHz range in at least one ear. If a subject met criteria in both ears, measurements were performed in each ear separately and averaged together to provide a single set of measurement per individual. Two groups of subjects who were specifically at risk for cochlear synaptopathy (CS) were tested; a middle-aged group (MA) with ages ranging from 36 – 60 years (N=58, 44 Female) and a young group with regular and substantial acoustic exposure (YExp) owing to membership in the Purdue Marching Band or a campus hunting/shooting club (N=53, ages 18 – 35 years, 27 Female). A young (ages 18 – 35) cohort of subjects who answered “No” to the question “Do you consider yourself regularly exposed to loud sound (e.g., play in a band, have a job with noisy tools/ machines)?” were included as the control group (YCtrl, N=55, 34 Female). The choice of the control group was conservative in that it was minimally restrictive, and likely includes subjects with a range of recreational acoustic-exposure histories that are representative of the local community in this age group. Note that our audiometric criteria for inclusion meant that a greater proportion of YExp and (an even greater proportion of) MA subjects who initially expressed interest had to be excluded compared to those in the YCtrl group.

Although our study design relied on using well-defined groups, a modified version of the noise exposure survey developed by Megerson (2010) was used to obtain a correlate of acoustic exposure in a subset of participants and establish whether the YCtrl and YExp groups were different in their reporting of exposures. The survey was modified to include two additional questions to ask participants about time spent in nightclubs and in bars/pubs as these are sources of noise exposure not queried on the original survey. Following the formulae provided in Megerson (2010), an average yearly noise exposure level was calculated for each subject. The results confirmed that the groups are indeed well-separated (Fig. S-1B).

### Chinchilla Cochlear Synaptopathy Model and Measurement Setup

Young (< 1.5 years old) male chinchillas (N=7) weighing 400 – 650 grams were used in accordance with protocols approved by the Purdue Animal Care and Use Committee (PACUC Protocol No: 1111000123). Awake and unrestrained chinchillas were exposed to 100 dB SPL, octave-band noise centered at 1 kHz for 2 hours. By systematically varying the exposure level, prior studies suggested that this exposure paradigm yields a temporary threshold shift (TTS) and concomitant broad cochlear synaptopathy (Hickox et al., 2017). The synaptopathic effects were histologically confirmed in the present study (see section on Inner Hair-Cell Synaptic Cochleogram). To further confirm TTS and characterize the effects of cochlear synaptopathy, a battery of non-invasive assays were measured from the chinchillas pre-exposure, 1-day post exposure, and 2-weeks post exposure (Fig. 1A). The non-invasive measures were acquired either with the animals anesthetized or awake, depending on the measure (described in separate sections on the individual assays), and were carried out in a double-walled, electrically shielded, sound-attenuating booth (Acoustic Systems, Austin, TX, USA).

For anesthetized measurements, anesthesia was induced with xylazine (4 mg/kg, subcutaneous), followed a few minutes later with ketamine (40 mg/kg, intraperitoneal). The xylazine reversal agent atipamezole (0.4 to 0.5 mg/kg, intraperitoneal) was used after procedures for faster recovery. Under anesthesia, eye ointment was used to keep the eyes lubricated and the animals’ vital signs were monitored throughout procedures with a pulse oximeter (Nonin 8600V, Plymouth, MN). An oxygen tube was placed near the animals’ nostrils. Body temperature was maintained near 37°C using a closed-loop heating pad with a rectal probe (50-7053F, Harvard Apparatus). Animals were provided lactated Ringer’s solution both before and after procedures (each 6 cc, subcutaneous).

For awake measurements, chinchillas were positioned in a cylindrical restraining tube (Fig. S-2, modified from Snyder and Salvi, 1994) that terminated in a neck-sized opening that was narrower than the rest of the tube. Movement was restricted in the axial direction of the tube by positioning the head and bullae outside the tube (i.e., beyond the narrow opening), and the rest of the body inside the tube. Head rotation was then restricted by placing a custom adjustable nose holder across the nose with the head at a comfortable downward angle that allowed for unrestricted airflow for breathing, and access to the ears for delivering auditory stimuli. A pulse oximeter (Nonin 8600V, Plymouth, MN) was placed on the pinna to monitor for any signs of restricted airflow, and a webcam was used continuously to monitor for any signs of discomfort. Animals remained comfortable throughout the ~30 minutes of awake data collection.

### Inner Hair-Cell Synaptic Cochleogram

The synaptopathy phenotype induced by our noise-exposure paradigm was confirmed by estimating synaptic loss in inner hair cells (IHCs) as a function of cochlear frequency location on a cohort of chinchillas (control: N=8; exposed: N=8) separate from the animals used for non-invasive physiological measures in this study. The synaptic loss was quantified by immunostaining and confocal imaging of CtBP2 (pre-synaptic ribbon protein). Animals were intracardially perfused with 4% paraformaldehyde in phosphate-buffered saline at pH 7.3. Cochleas were dissected and immediately perfused through the cochlear scalae, post-fixed for 2 h at room temperature, dissected into six pieces without decalcification (roughly half turns of the cochlear spiral) for whole-mount processing of the cochlear epithelium. Cochlear pieces were transferred to net well sieve in 5 ml disposable cup with ~1 cc of 30% sucrose and freeze-thawed for membrane permeabilization (Liberman and Liberman, 2015). Immunostaining began with a blocking buffer (PBS with 5% normal horse serum and 0.3—1% Triton X-100) for 1 h at room temperature and followed by overnight incubation at 37 °C with mouse (IgG1) anti-CtBP2 (C-terminal Binding Protein), to quantify pre-synaptic ribbons and rabbit anti-Myosin VIIa to delineate the IHC cytoplasm. Primary incubations were followed by two sequential 60-min incubations at 37 °C in species appropriate secondary antibodies (coupled to Alexafluor dyes) with 0.3—1% Triton X-100. To understand the position of synapses with respect to IHC surface, anti-myosin VIIa & Topro-3 (nuclei dye) were used. Confocal z stacks of frequency-specific regions from each ear were obtained using a high resolution (1.4 NA) oil immersion objective (63x) on an inverted Zeiss LSM 710. Images were acquired in a 1448 × 1604 raster (pixel size = 0.036 *μ*m in x and y, scan speed 7; Averaging of 2 with 8 bit depth) with a z-step-size of 0.25 mm. Image stacks (.lsm files) were ported to an off-line processing station, where further 3-D morphometry was performed using a commercial image-processing program (FEI’s Amira, Visage Imaging). Synaptic ribbons were counted and divided by the total number of IHC nuclei in the microscopic field including fractional estimates. The percentage of IHC synapse loss was plotted as a function of cochlear frequency location (Fig. 1B).

### Behavioral Audiometry (Humans only)

Standard clinical and extended-high-frequency audiograms were measured using a GSI AudioStar Pro audiometer (Eden Prairie, MN) and Sennheiser HDA 200 high-frequency headphones by employing pulsed-tone stimuli. Thresholds were determined for 0.25, 0.5,1, 2, 3, 4, 6, 8, 10, 12.5, 14 and 16 kHz using a bracketing procedure (modified Hughson and Westlake procedure). An important feature of this study was that the three human groups (YCtrl, YExp, and MA) were tightly matched in audiometric thresholds with mean and median differences being < 5 dB at every frequency tested from 0.25 and 8 kHz (Fig. S-1A). The YCtrl and YExp groups were also matched in the 8 – 16 kHz range (with mean and median differences < 5 dB, again). Owing to substantial effect of age on hearing sensitivity in the extended high-frequency range (Lee et al., 2012), it was infeasible to match the young and MA groups in this range. The MA group thus showed an approximately 25-dB threshold elevation in the extended-high-frequency range. When performing statistical analyses of putative assays of cochlear synaptopathy, the audiograms were summarized into three bins: a low-frequency average (LFA; 0.25 – 3 kHz), a high-frequency average (HFA; 4 – 8 kHz), and an extended-high-frequency average (EHFA; 10 – 16 kHz). The mean and median differences were < 5 dB across all three groups for LFA and HFA, and also across YCtrl vs. YExp for EHFA. This tight matching in audiograms between the groups allowed for interpreting the other suprathreshold measures in the context of cochlear synaptopathy.

### Wideband Middle-Ear Muscle Reflex (WB-MEMR)

Motivated by findings in mice (Valero et al., 2018), the WB-MEMR was a key candidate for an assay of cochlear synaptopathy and was measured in both species using shared methods and software. Chinchilla measurements were acquired with the animals awake and restrained. Human subjects passively watched a muted video with subtitles during data acquisition. The WB-MEMR paradigm was adapted from (Keefe et al., 2017). An ER-10X wideband measurement system was used to acquire data from human subjects whereas an ER-10B+ system coupled to ER-2 transducers and foam eartips were used for chinchilla data collection (Etymotic, Elk Grove Village, IL). Both measurement systems allowed for probe stimuli and ipsilateral reflex-eliciting stimuli to be presented from separate speakers to limit interchannel interactions and distortions, and consisted of a microphone to measure sound pressure near the ear tips. A 90-dB peSPL click probe with a flat incident power spectrum in the 0 – 10 kHz range was used to measure the acoustic immittance properties of the ear-canal middle-ear system. Each WB-MEMR measurement trial consisted of a series of seven clicks alternating with 120 ms-long ipsilateral noise elicitors (Fig. S-3A). The gap between the peak of the click and the onset ramp of the noise elicitor was 27.94 ms, whereas the gap between the offset ramp of the noise and the next click was 13.97 ms. This trial structure was used for each elicitor level and the level series was repeated 32 times with an intertrial interval of 1.5 seconds to allow for the middle-ear immittance to relax back to baseline levels. For each elicitor level, the immittance measured using clicks numbered two through seven in the sequence were averaged together and the change relative to the first click was calculated as the WB-MEMR metric. Although the data reported in the present study represent averaged measurements from 32 trials, the MEMR was measurable with single trials and thus is readily translatable to clinical applications with limited time availability. The immittance change from the WB-MEMR protocol was quantified using analogous metrics for chinchillas and humans. For both species, the dB-change in ear-canal pressure induced by the MEMR was quantified as a function of frequency to yield a pattern of alternating negative and positive peaks at different frequencies (Fig. S-3B, Fig. S-4A). For chinchillas, the absolute value of this pattern was averaged over 0.5 – 2 kHz to yield a single number per elicitor level (Fig. S-4B). For human data, additional calibrations which help reduce extraneous variance were leveraged (as described in the Acoustic Calibrations section). Accordingly, the dB-change in ear-canal pressure was added to the dB-change in ear-canal conductance (Fig. S-3C) to yield a stereotypical pattern of dB-change in absorbed power (Keefe et al., 2017, Fig. S-3D). The absolute value of this dB-change in absorbed power was averaged over 0.5 - 2 kHz to yield a single number per elicitor level (Fig. S-3E). Chinchilla WB-MEMR data were acquired for broadband (BB) noise (0.5 – 8 kHz) elicitors in the 34 – 94 dB SPL range in steps of 6 dB. Human WB-MEMR data were acquired both for BB (0.5 – 8 kHz) and highpass (HP, 3 – 8 kHz) elicitors in the 34 – 88 dB FPL (Forward-pressure level) range in 6 dB steps. Noise spectra had a logarithmic roll-off with frequency in both species to produce a flat excitation pattern including adjustments for the sharpening of cochlear tuning at higher frequencies (Shera et al., 2002). A thresholding procedure was used to reject artifactual trials before averaging. The WB-MEMR growth functions (with elicitor level) were baseline corrected by subtracting the mean values across the 34 and 40 dB SPL/FPL points. For plotting and group level analysis, a three-parameter sigmoid function was fit to individual baseline-adjusted WB-MEMR growth functions. The level at which the fitted function crossed 0.1 dB was recorded as the WB-MEMR threshold for each individual.

### Acoustic Calibrations to Reduce Human Ear-Canal Filtering Effects

For human measures, the ER-10X probe was calibrated using classic methods for characterizing two-port networks (Allen, 1986). The Thevenin-equivalent source pressure and impedance for the click probe were estimated by measuring the acoustic response at the ER-10X microphone when the eartip was coupled to loads whose acoustic impedance values can be approximated using theoretical calculations (brass cylinders of 8 mm diameter and five different lengths, that is the ER-10X calibrator). The estimates were refined until the so-called “calibration error” (a dimensionless energy ratio scaled by a factor of 10000, and averaged over 2–8 kHz) was minimized. Error values of < 1 are typically considered good quality calibration (Neely and Liu, 1994); we routinely obtained errors in the 0.01 – 0.04 range. With the probe properties calibrated, the same click stimulus was then used to estimate the immittance properties of each subject’s ear for each insertion of the probe tip. Following Groon et al. (2015), we considered the probe as adequately sealed in the ear canal if the low-frequency (0.2 - 0.5 kHz average) ear absorbance was less than 29% and the admittance phase was greater than 44°. The in-ear calibration measurements were used to derive a voltage to forward-pressure-level (FPL) transfer function that was then used to generate voltage waveforms that would yield stimuli at desired FPL. These calibration methods help reduce extraneous variability from ear-canal filtering both within and across individuals (Bharadwaj et al., 2019).

### Distortion-Product Otoacoustic Emissions (DPOAEs)

To indirectly assess cochlear mechanical function and outer-hair-cell (OHC) integrity, DPOAEs were measured in both species using logarithmically sweeping primaries (Long et al., 2008). f2 frequency was swept down from 16 kHz to 2 kHz with f1 = f2/1.225. For human measurements, because the additional calibrations were available (as described in the Acoustic Calibrations section), the primary levels were kept constant across frequencies at 66/56 dB FPL. The use of constant-FPL primaries allowed for the 2f1 - f2 DPOAE level to be quantified in emitted-pressure level (EPL) units (Charaziak and Shera, 2017). For chinchilla measurements, constant-SPL primaries at 75/65 dB SPL were used instead with the DPOAE also quantified in SPL units. The identical least-squares approach with 0.5 s-long windows was used in both species to extract the DPOAE; this choice of window length was guided by previous work suggesting that this allows for extracting the contribution of just the distortion source (Abdala et al., 2015). To cross-validate our swept-tone methods, standard DP-grams were also acquired from the chinchillas using pure-tone primaries. The DPOAEs acquired using pure tones were comparable to those acquired using swept-tone primaries (Compare Fig. S-5, and Fig. 1C).

### Auditory Brainstem Responses (ABRs)

ABRs were measured in human subjects using a 32-channel EEG cap (Biosemi, Amsterdam, Netherlands) using HP (3 – 8 kHz), 105 dB SPL click stimuli and in-ear gold-foil tiptrodes (ER3-26A combined with ER3-28S) coupled to earphone transducers (ER2, Etymotic Research, Elk Grove Village, IL). The use of HP clicks helps reduce extraneous inter-subject variability owing to cochlear dispersion, while simultaneously focusing the assay on the basal regions where audiometric threshold shift owing to noise-induced hearing loss tend to appear (Bharadwaj et al., 2019). The presentation timing of the clicks was random with a Poisson distribution (Polonenko and Maddox, 2019) with a mean rate of 11 clicks per second and randomized polarity with 4000 repetitions per polarity. The ABR was extracted using conventional trial averaging. To quantify the amplitudes of wave I and wave V without the experimenter bias on group differences, an automated procedure based on dynamic time warping was used (Picton et al., 1988). The grand-averaged waveform across all subjects (across all three groups) was defined as the template ABR. Wave I and V peaks and troughs were marked manually on this template waveform; the corresponding points identified by automatic dynamic time warping were identified as the ABR wave I and V peaks and following troughs for individual subjects. The peak-to-trough swings were then extracted as the wave I and V amplitudes. Wave I/V ratio formed the second candidate assay for cochlear synaptopathy (Bharadwaj et al., 2019). Separate wave I and V values are plotted in Fig. S-6.

Anesthetized ABRs in response to tone-burst stimuli were measured from chinchillas pre-TTS exposure and 2-weeks post-TTS exposure. Measurements were acquired using subdermal needle electrodes with a stimulus presentation rate of 20 bursts per second and as a function of level, with 500 repetitions of both positive and negative polarities collected and averaged. ABR thresholds were computed using our standard cross-correlation technique (Henry et al., 2011) at the two time points and were used to establish that the exposure did not lead to a significant threshold shift (i.e., < 5 dB; Fig. 1D). Suprathreshold amplitudes of the ABRs (averaged at 60, 70, and 80 dB SPL) were averaged across the 4 and 8 kHz tone-burst conditions to yield a metric that is analogous to the high-level 3 – 8 kHz clicks used in humans. The ABR wave I/V ratio was again evaluated as a candidate metric for cochlear synaptopathy. Separate wave I and V values are shown in Fig. S-7.

### “Clinic-Style” Measures in Humans

To compare the sensitivity of ABR and MEMR protocols available in clinical systems to the sensitivity of lab-based targeted procedures, the same human participants were also measured using standard clinical equipment. ABRs were measured using a SmartEP system (Intelligent Hearing Systems, Miami, FL) in response to 80 dB nHL clicks at a presentation rate of 11.1 clicks per second, a gain setting of 150k, and a 50-3000 Hz bandpass filter. ABR wave I and V peaks and troughs were manually marked by trained research assistants who were also graduate students in the clinical audiology program at Purdue University. MEMRs were measured using a Titan System (Interacoustics, Assens, Denmark) in response to 4-kHz tone elictors between 75 and 100 dB HL in 5 dB steps. A 226 Hz tone probe was used for MEMR measurements, as is customary for adult immittance measurements in audiology clinics.

### NHANES 2011–2012 MEMR Data

A large repository of human audiological data is available from the Centers for Disease Control and Prevention (CDC) as part of the National Health and Nutrition Examination Survey (NHANES). This repository includes MEMR data acquired using the Earscan system and audiometric data in the standard clinical frequency range. MEMR-induced immittance change waveforms (in “Earscan” units) in response to two presentations of a 105 dB HL 2 kHz tone elicitor are available from 4500 adult subjects in the 20 - 69 year age range. Given the results from lab-based WB-MEMR measures suggesting that middle-aged individuals exhibit cochlear nerve degeneration across the frequency range, we asked whether there is evidence of such degeneration in this publicly available dataset. To minimize the contribution of audiometric hearing loss to the MEMR results, we only included the subset of N=1884 subjects whose audiometric thresholds at 8 kHz was 20 dB HL or better and age of 60 years or less. Following Flamme et al. (2017), the MEMR traces were expressed in Z-scores relative to the last 450 ms of the available recordings. The MEMR amplitude was quantified as the mean Z-score between 350 and 450 ms after the onset of the 2 kHz tone elicitors. This value was plotted as a function of age in 2-year bins, with each containing about 100 subjects (Fig. 3B) and revealed a steady decline with age despite clinically normal hearing. For effect-size calculations the participants were divided into two groups matching the definition of the participant groups recruited in the present study.

### Statistical Analysis

Statistical inference was performed by fitting linear mixed-effects models (Box and Tiao, 2011) to the data when multiple data points were measured for the same individual (WB-MEMR data in both species, and pre vs. 2-week-post chinchilla ABR data), or by fitting simple linear models when only one data point was available per subject (Human ABR data). Fixed-effects terms were included to model the effects of group (humans), time-point (pre vs. 2-week-post in chinchillas), and the effects of various covariates (sex, audiometric thresholds, DPOAE amplitudes in humans), whereas subject-related effects were treated as random effects. Homoscedasticity of subject-related random effects was not assumed initially and hence the error terms were allowed to vary and be correlated across the levels of fixed-effects factors. In order to not over-parameterize the random effects, the random terms were pruned by comparing models with and without each term using the Akaike information criterion and log-likelihood ratios (Pinheiro and Bates, 2000). The best fitting random-effects model turned out to be a single subject-specific random effect that was condition independent in all cases. This random-effect term was used for all subsequent analysis. All model coefficients and covariance parameters were estimated using restricted maximum likelihood as implemented in the lme4 library in R (Bates et al., 2007). To make inferences about the experimental fixed effects, the F approximation for the scaled type-II Wald statistic was employed (Kenward and Roger, 1997). This approximation is more conservative in estimating false-alarm rates than the Chi-squared approximation of the log-likelihood ratios and has been shown to perform well even with fairly complex covariance structures and small sample sizes (Schaalje et al., 2002). The p-values and F-statistics based on this approximation are reported. For the human ABR data, because a simple linear model is used, group effects are evaluated using T-statistics.

Beyond formal statistical inference on our two key lab-based assays, effect size calculations were performed for each individual lab-based and clinic-style measure using the bias-corrected procedures outlined in Hedges (1982). Given the larger human sample sizes, mean and standard deviation for effect size metrics were computed using median-based estimates. For the chinchilla data, given smaller sample size, conventional sample statistics were used.

### Blind Classification of Human Groups

To assess the sensitivity of our battery of lab-based measures as a whole (rather than individual measures) in predicting acoustic exposure group status and middle-age status, we used a support-vector machine (SVM) classifier to blindly classify individuals into their corresponding groups. WB-MEMR thresholds, high-level amplitudes for both HP and BB elicitors, and the ABR wave I/V ratio were all entered as features (five total) for the classifier to operate on. To obtain the mean and standard error of the classification accuracy, a leave-1-out train-test split procedure was used. The resulting classification accuracies are reported in Fig. 3C. The equivalent effect sizes for the hybrid metric learned by the classifier were estimated from the classification accuracy by computing the p-value of the classifier against a binomial null distribution of chance classification (50%) and using the p-value to obtain equivalent Cohen’s d scores (Rosenthal and Rubin, 2003). These combined/hybrid effect sizes are reported in Fig. 3A.

## Supplementary Figures

**Figure S-1.**
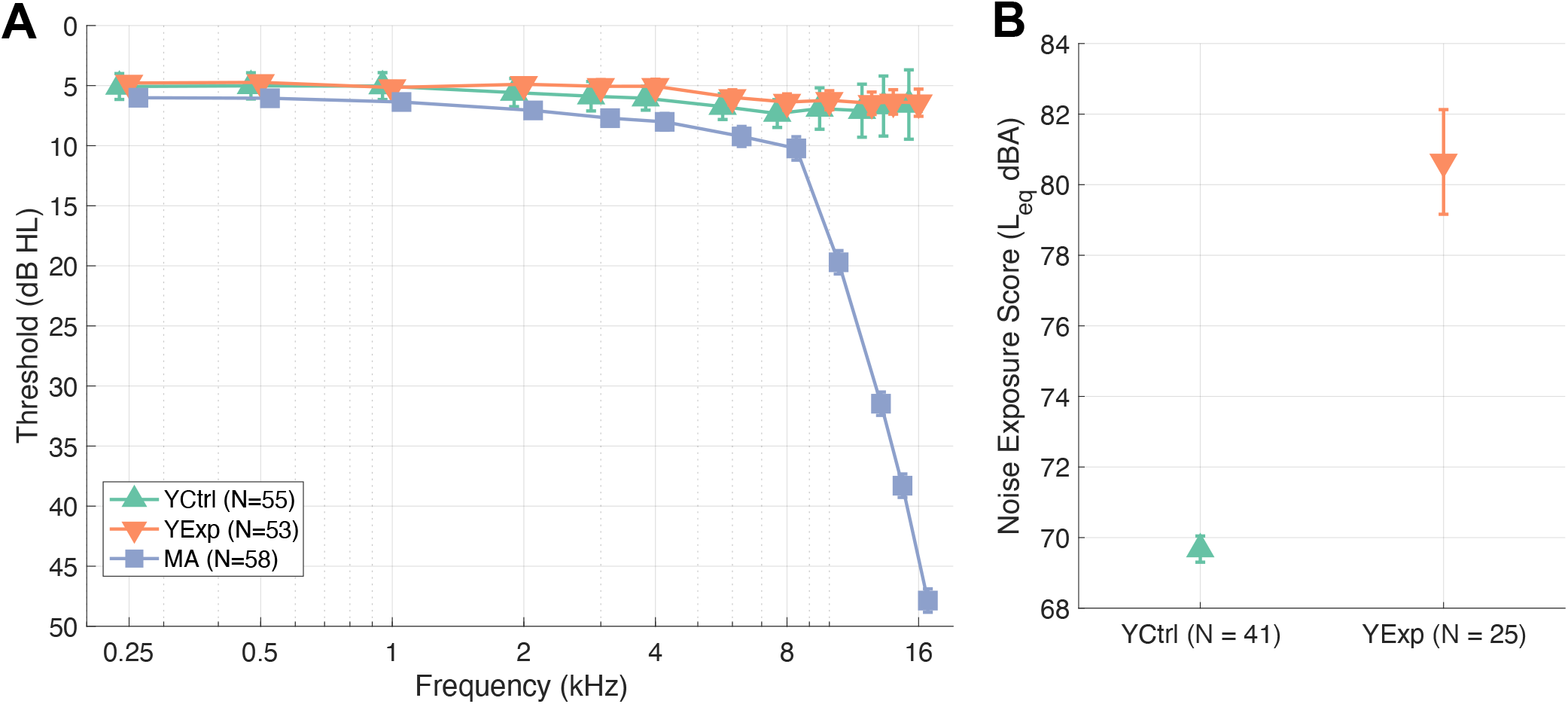
(A) Audiometric thresholds show tight matching between groups in the standard clinical frequency range (up to 8 kHz). YCtrl and YExp groups are tightly matched up to 16 kHz. (B) As expected, noise-exposure scores from a subset of young controls (YCtrl) and young noise-exposed subjects (YExp) using the approach in Megerson (2010) confirms that the two are well-separated in their reporting of acoustic exposures. Data points represent mean ± STE bars.

**Figure S-2.**
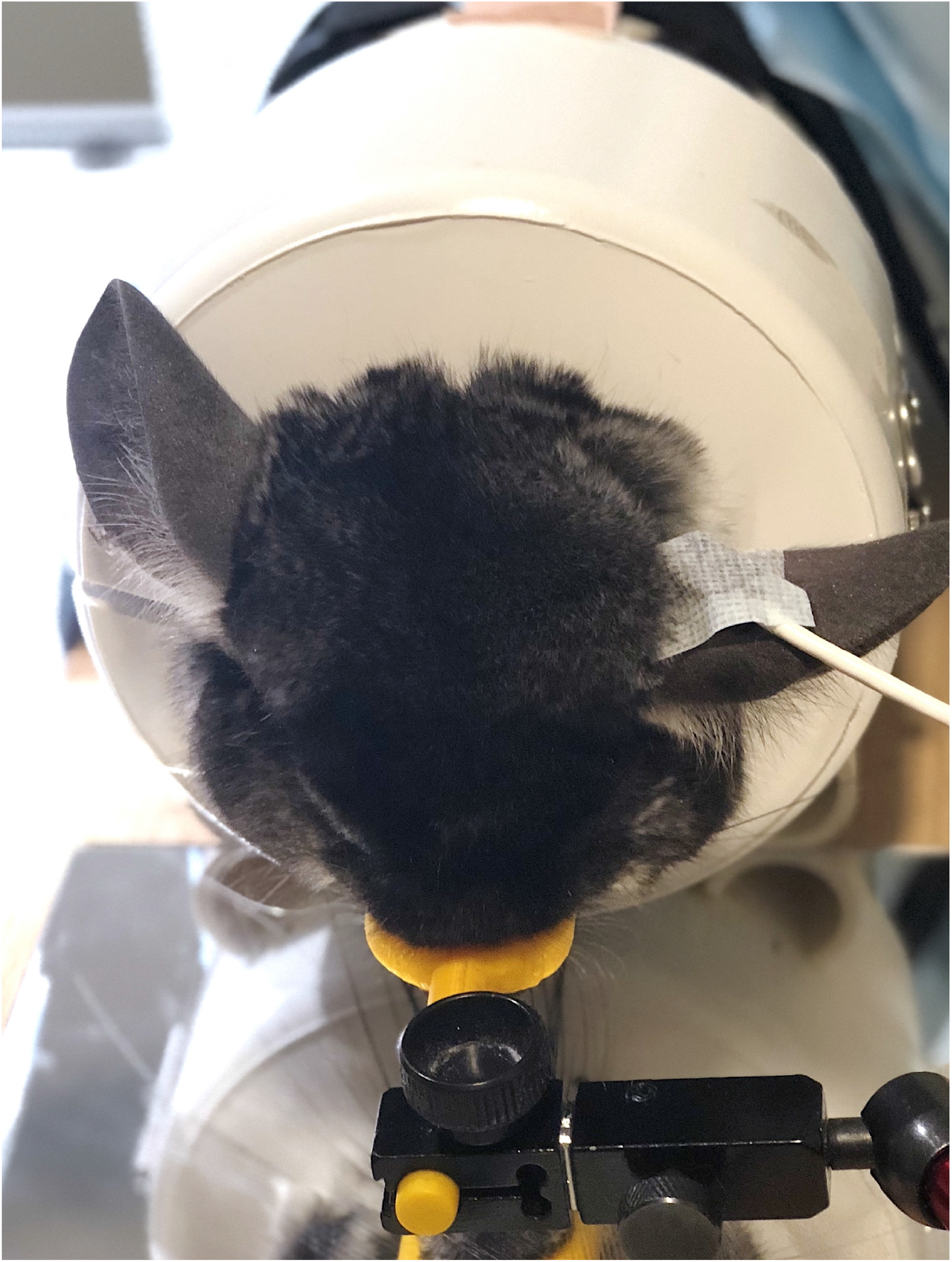
Photograph of awake chinchilla setup for DPOAE and WB-MEMR measurements.

**Figure S-3.**
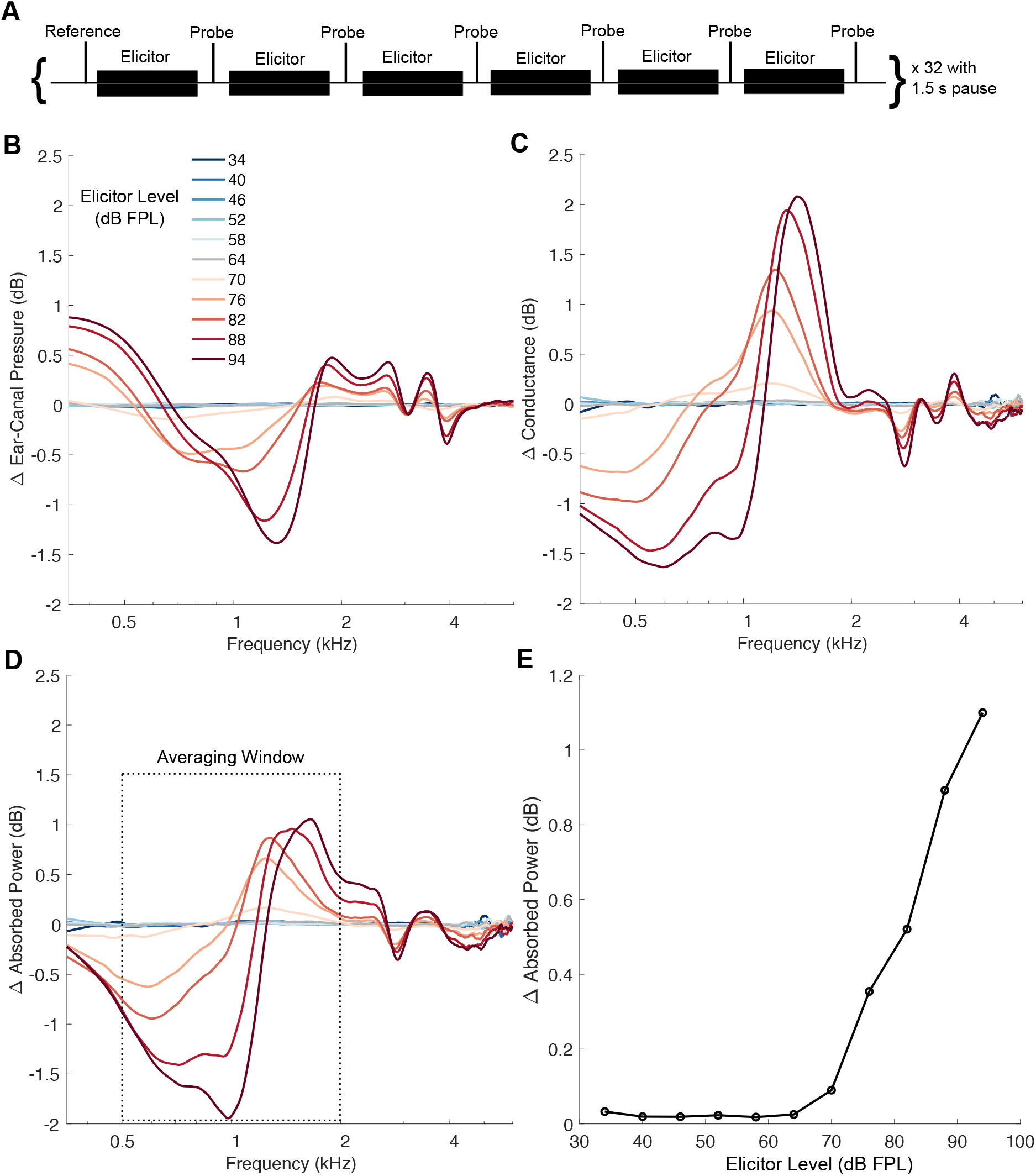
WB-MEMR measurement paradigm and example data from a human participant. (A) Trial sequence consisting of alternate click probe and ipsilateral elicitors. (B) Frequency pattern of WB-MEMR immittance change quantified as change in ear-canal pressure for the probe click. (C) Conductance change associated with MEMR. (D) WB-MEMR immittance change quantified as change in absorbed power (sum of values in B and C). (E) Raw (before baseline correction and sigmoid curve fitting) WB-MEMR growth function with elicitor level extracted by averaging absolute dB-changes in absorbed power in the 0.5–2 kHz range.

**Figure S-4.**
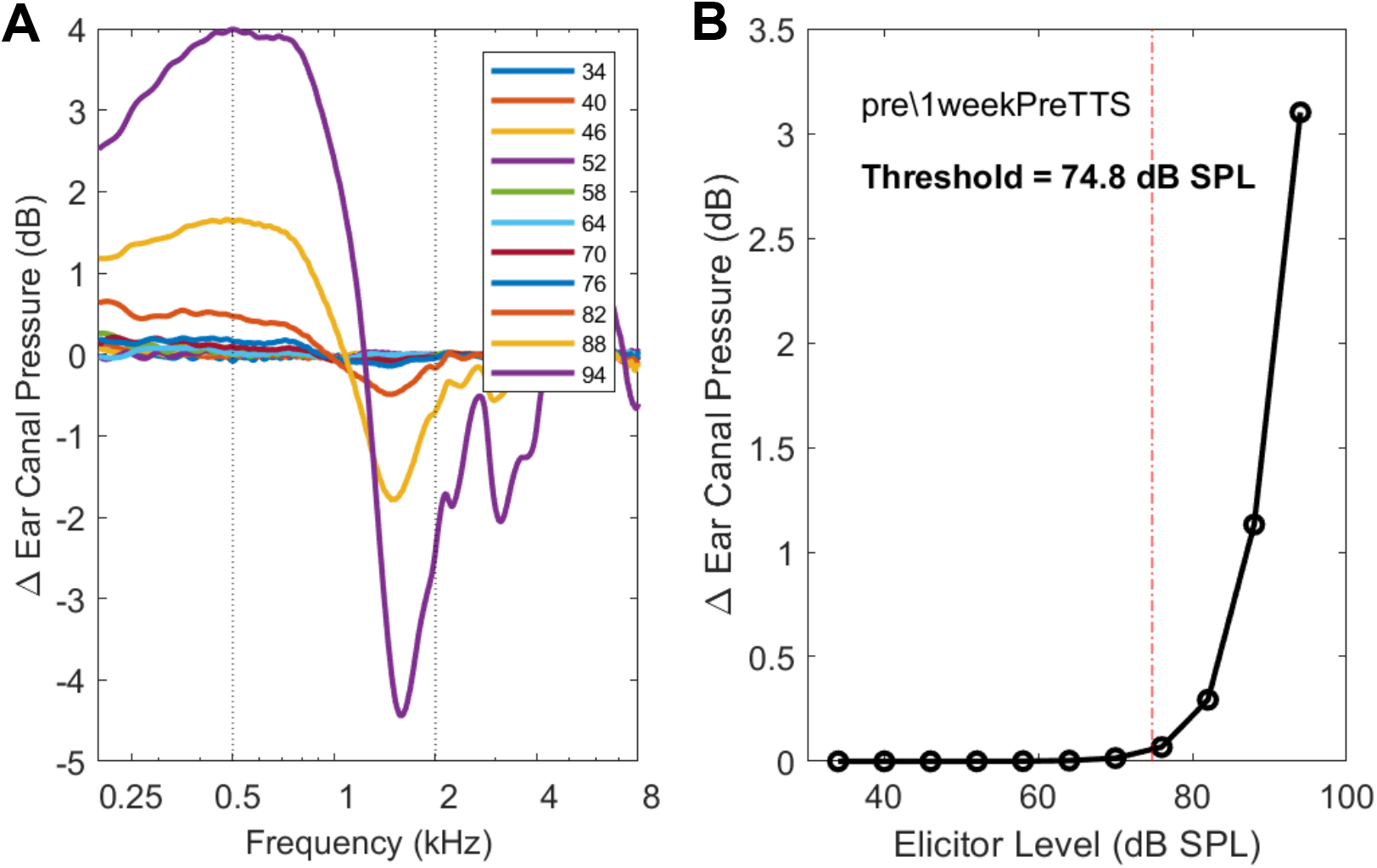
WB-MEMR example from a chinchilla participant. (A) Stereotypical alternating profile of WB-MEMR response across frequency. (B) Fitted WB-MEMR growth function with elicitor level extracted by averaging absolute dB-changes in ear-canal pressure in the 0.5 – 2 kHz range and fitting a sigmoid growth function.

**Figure S-5.**
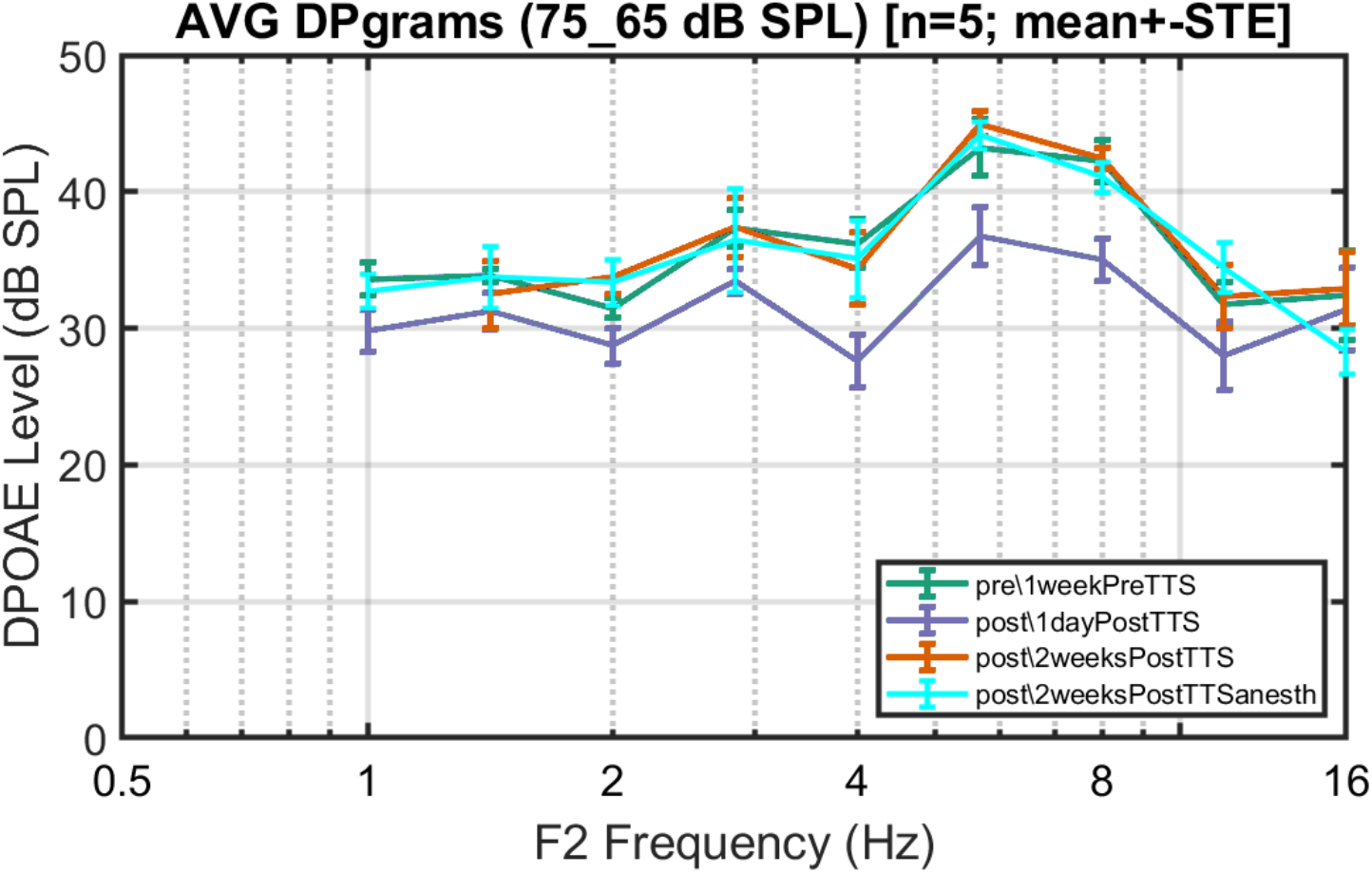
DPOAEs measured using pure-tone primaries instead of the swept-tone protocol. Response values are comparable to swept-tone measurements in Fig. 1C. Data points represent mean ± STE bars.

**Figure S-6.**
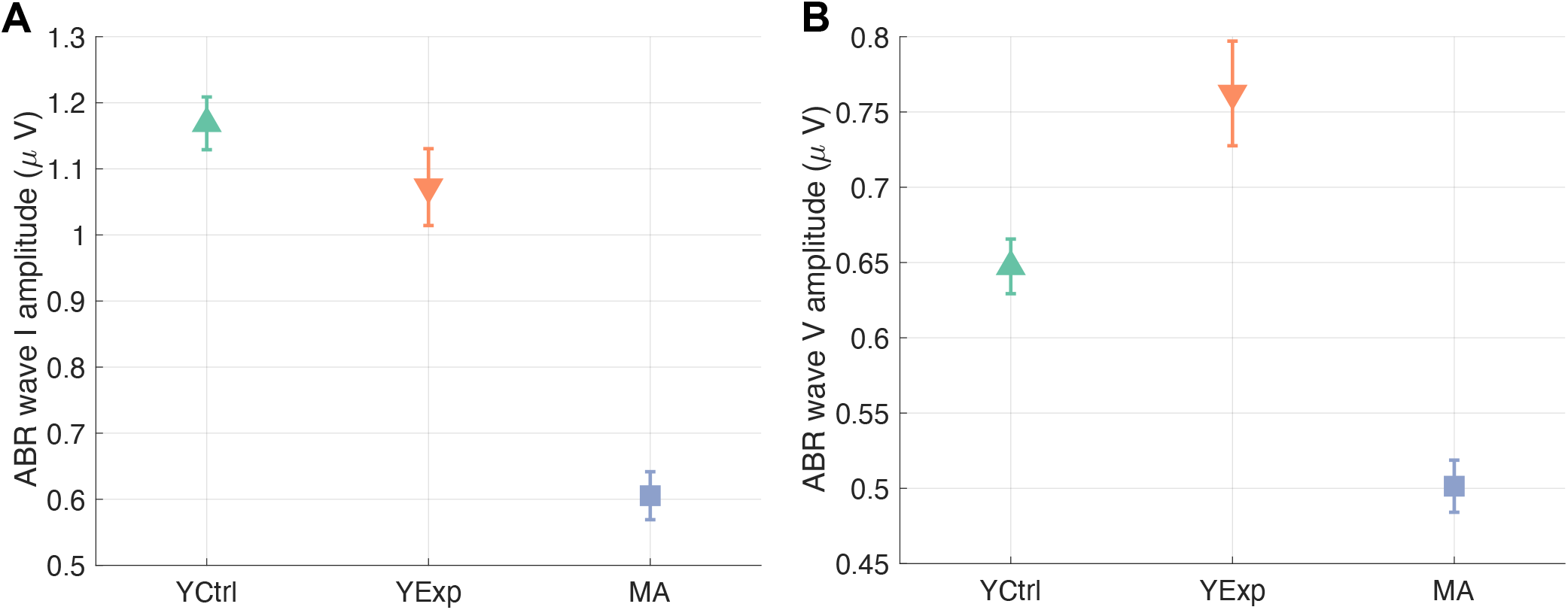
Separate wave I (A) and wave V (B) values shown for the three human groups. The ABR wave I/V ratio derived from these values was used as a putative assay of cochlear synaptopathy (Fig. 2C). Data points represent mean ± STE bars.

**Figure S-7.**
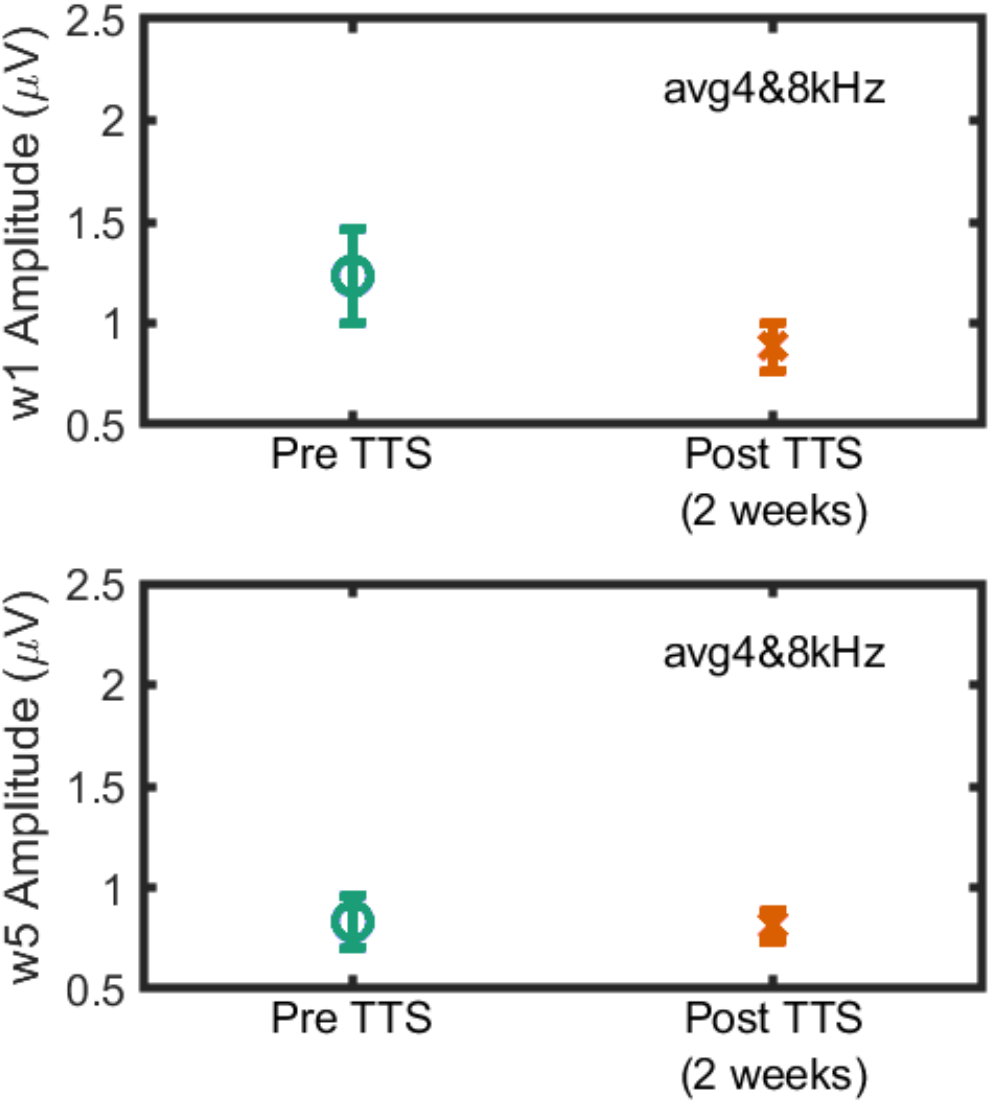
Separate wave I (top) and wave V (bottom) values shown for chinchilla subjects. The ABR wave I/V ratio derived from these values was used as a putative assay of cochlear synaptopathy (Fig. 1F). Data points represent mean ± STE bars.

## Footnotes

1 Full DOI will be provided upon peer-reviewed publication

